# Discovery of an anti-virulence compound that targets the *Staphylococcus aureus* SaeRS two-component system to inhibit toxic shock syndrome toxin 1 (TSST-1) production

**DOI:** 10.1101/2024.02.27.582338

**Authors:** Karine Dufresne, Dennis A. DiMaggio, Carla S. Maduta, Shaun R. Brinsmade, John K. McCormick

## Abstract

Menstrual toxic shock syndrome (mTSS) is a rare but severe disorder associated with the use of menstrual products such as high-absorbency tampons and is caused by *Staphylococcus aureus* strains that produce the toxic shock syndrome toxin-1 (TSST-1) superantigen. Herein, we screened a library of 3920 small bioactive molecules for the ability to inhibit transcription of the TSST-1 gene without inhibiting growth of *S. aureus*. The dominant positive regulator of TSST-1 is the SaeRS two-component system (TCS), and we identified phenazopyridine hydrochloride (PP-HCl) that repressed production of TSST-1 by inhibiting the kinase function of SaeS. PP-HCl competed with ATP for binding of the kinase SaeS leading to decreased phosphorylation of SaeR and reduced expression of TSST-1 as well as several other secreted virulence factors known to be regulated by SaeRS. PP-HCl targets virulence of *S. aureus*, but it also decreases the impact of TSST-1 on human lymphocytes without affecting the healthy vaginal microbiota. Our findings demonstrate the promising potential of PP-HCl as a therapeutic strategy against mTSS.

## Introduction

Menstrual toxic shock syndrome (mTSS) is a serious bacterial toxin-mediated disease that became widely recognized in the early 1980s due to an epidemic of cases in the United States that was associated with the use of high-absorbency tampons (1). Following early investigations into this epidemic, it was determined that these women were vaginally colonized with *Staphylococcus aureus* that produced a unique toxin now known as toxic shock syndrome toxin-1 (TSST-1) (2, 3). TSST-1 functions as a bacterial superantigen, forcing the activation of numerous T cells that can progress to a cytokine storm that characterizes mTSS (4). Approximately 30% of the general human population is understood to be colonized by *S. aureus* within the nasal passages (5). However, many other body sites are also frequently colonized (6) and in healthy women using tampons, ∼30-40% were vaginally colonized with *S. aureus*, with ∼5% of strains producing TSST-1 (7, 8). The use of high-absorbency tampons is understood to have altered the vaginal environment to create conditions that are sensed by *S. aureus* which can result in the increased production of TSST-1 (9–12). Although tampons have been re-designed to reduce the risk of mTSS, the importance of mTSS for public safety remains, and understanding the regulatory pathways that control TSST-1 production in conditions mimicking the vaginal tract may be key for prevention of future cases of mTSS.

The incidence of mTSS in the United States is ∼0.5 to 1.0 per 100,000 population (13), which is far below the number of women who are vaginally colonized by TSST-1^+^ *S. aureus*. Thus, multiple factors are likely protective for the development of mTSS including the proper use of menstrual management products (14), the presence of neutralizing anti-TSST-1 antibodies (15), the endogenous vaginal microbiota (16–19), and environmental signals that repress TSST-1 production including low levels of O_2_ and CO_2_ levels (20), acidic pH (21), and high levels of glucose (12). Furthermore, *S. aureus* colonization will also vary during the menstrual cycle as the environment is dynamic due to hormonal fluctuations (12, 22).

As an alternative to the use of antibiotics with bacteriostatic or bactericidal activity, genetic regulatory systems in *S. aureus* have been targeted for anti-virulence therapeutics. For example, the accessory gene regulator (Agr) two-component system (TCS) is a well-studied quorum sensing system that is activated via the production of endogenous auto-inducing peptides (AIPs), and apart from SaeRS, Agr is the other major exotoxin regulator in *S. aureus* (23). Indeed, Agr controls the expression of TSST-1 by de-repression of the Rot repressor (24). Coagulase negative staphylococcal (CoNS) species, as well as *S. aureus*, can produce AIP variants that can function to inhibit heterologous Agr systems. For example, interference with Agr signaling via inhibitory AIP molecules produced by *Staphylococcus hominis* could suppress skin damage and inflammation in a mouse model of *S. aureus*-induced atopic dermatitis (25) and reduce lesion sizes in a mouse model of dermonecrosis without altering bacterial counts (26). However, targeting the Agr system could also potentially evolve strains into a persistent *agr*-deficient state (27).

The GraXRS sensing system is another TCS that responds to cell-envelope stress including cationic antimicrobial peptides and low pH (28, 29). A small molecule screen designed to find inhibitors of the early step in wall teichoic acid production identified a GraR inhibitor, which blocked intracellular survival of *S. aureus* within macrophages and enhanced larvae survival in a *Galleria mellonella* infection model (30). Furthermore, the autolysis-related locus TCS (ArlRS) controls many phenotypes including adhesion, capsule production and metal transport, primarily through the MgrA stand-alone transcription factor (31). Using a *mgrA* promoter reporter, small molecule inhibitors of ArlRS could reduce skin infections, but did not alter viable *S. aureus* recovered from the lesions (32).

Recently, an inhibitor of SaeS was found using an α-hemolysin promoter screen that inhibited SaeRS regulated virulence factors, but also appeared to affect Agr, and was able to constrain experimental invasive infections by *S. aureus* (33). A structure-based virtual screen also identified the non-steroidal anti-inflammatory drug Fenoprofen as a direct SaeR inhibitor that could attenuate *S*. *aureus* virulence *in vitro* and *in vivo* (34). Although these compounds have not yet reached therapeutic use in humans, there are now multiple examples by which anti-virulence compounds demonstrate the potential of targeting *S. aureus* TCSs to disrupt virulence and potentially to increase the efficiency of antibiotic therapy.

To advance therapeutics that could further inhibit the production of TSST-1, we developed a cell-based platform using a TSST-1 transcriptional luciferase reporter assay with *S. aureus* grown in a vaginal-mimicking media to simulate the environmental conditions of mTSS. Using this platform, we screened a library of small bio-active molecules and identified that phenazopyridine hydrochloride (PP-HCl) demonstrates anti-virulence activity against TSST-1 expression without growth inhibitory properties. This molecule was further characterized to decipher its anti-virulent mechanisms, and we demonstrate that PP-HCl functions through inhibition of kinase activity of the SaeRS TCS, the major positive transcriptional activator of TSST-1 (35).

## Results

### PP-HCl represses production of TSST-1 without inhibiting S. aureus growth

We screened a library of 3920 bioactive molecules (**Fig. 1**), each at 10 µM, for the ability to repress the activity of the TSST-1 promoter (P*_tst_*) using *S. aureus* MN8 harboring pAmilux::P*_tst_*, a plasmid that indirectly measures P*_tst_* activity using a luciferase (*lux*) reporter (**Table 1**). Compound screening was performed in vaginally-defined media (VDM) containing low amounts of glucose (700 μM) to mimic the vaginal environment during mTSS when CcpA-mediated repression of TSST-1 is relieved (12). From the initial screen, seventy molecules demonstrated low luminescence (less than 100 raw RLU) with growth equal or greater than 95% of the control cultures (**Fig.1**). Among these compounds, 18 were previously described to possess antimicrobial activity and were not followed further (**Fig.1)**. The remaining 52 hits were randomly tested under low throughput conditions and titrated using the *S. aureus* P*_tst_* reporter strain. In this context, PP-HCl showed no growth inhibition (**Fig.2A**) with robust repression of P*_tst_* with 5µM (**Fig.2B and 2C**). To corroborate the luciferase reporter experiments, TSST-1 protein levels from *S. aureus* MN8 supernatants were evaluated using an anti-TSST-1 Western Blot. A similar trend was observed with decreased TSST-1 produced when *S. aureus* was grown in 5 μM PP-HCl with only a faint band visible when grown in 50 μM (**Fig.2D**). These data demonstrate that PP-HCl decreases *tst* promoter activity and the production of TSST-1 protein without inhibiting *S. aureus* growth and may represent a new potential anti-virulent compound for *S. aureus*.

**Figure 1.**
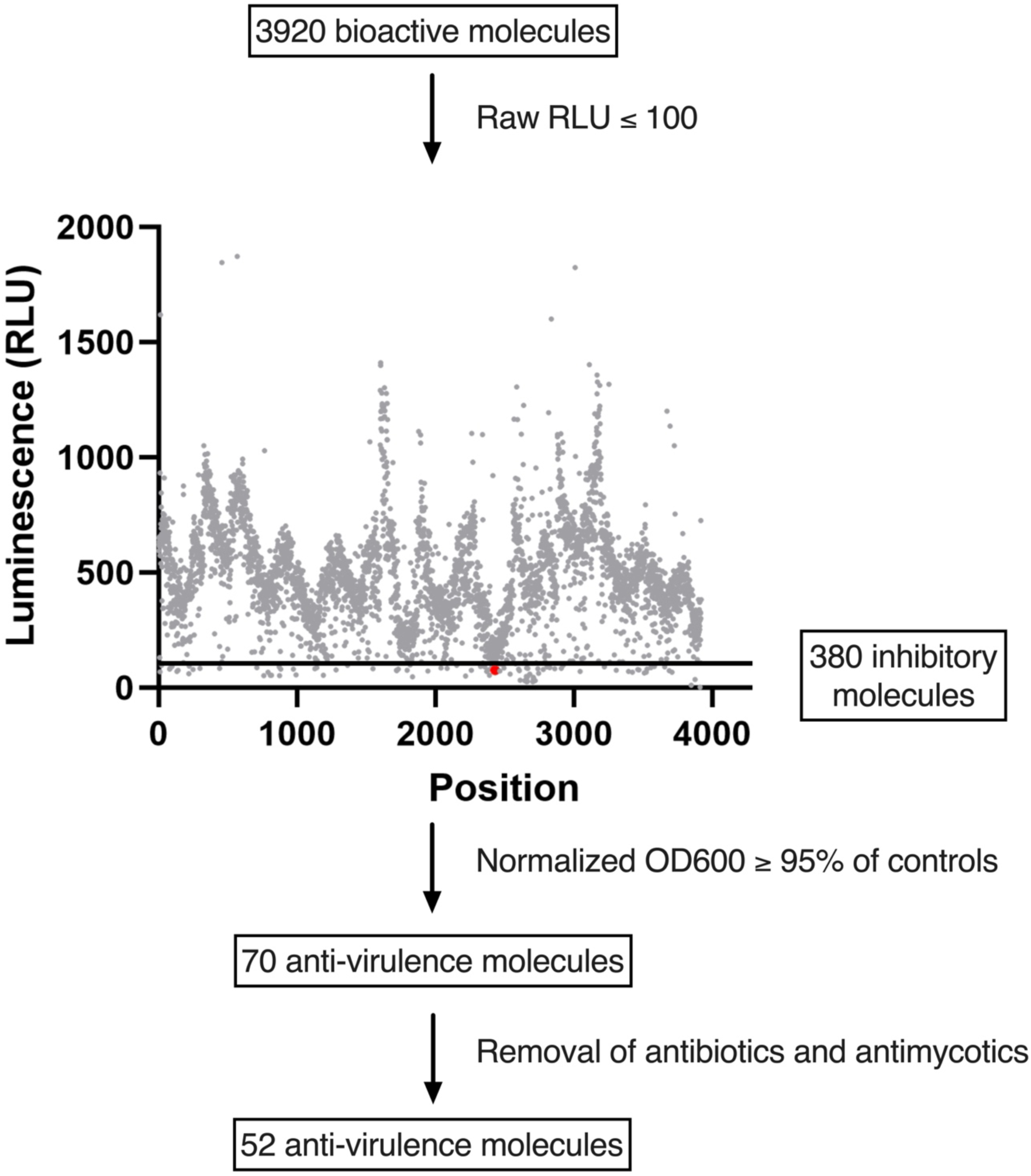
Screening strategy for anti-virulent against TSST-1. *S. aureus* MN8 reporter assay monitoring growth and luminescence production (expression of the promoter P*_tst_*) was performed with 3920 bioactive molecules each tested at 10µM concentration. The graph represents all compounds tested with the 100 RLU threshold represented by an additional line; PP-HCl is marked in red. Compounds with a raw luminescence equal or less than 100 RLU were further selected (380 molecules). Compounds lacking antimicrobial activity with normalized OD_600_ equal or greater than 95% of the control were selected leading to 70 putative inhibitory molecules. An additional selection was made for the bioactive molecules with no known previous antimicrobial activity (both antibiotics and antimycotics) resulting in 52 molecules of interest from the initial screen.

**Figure 2.**
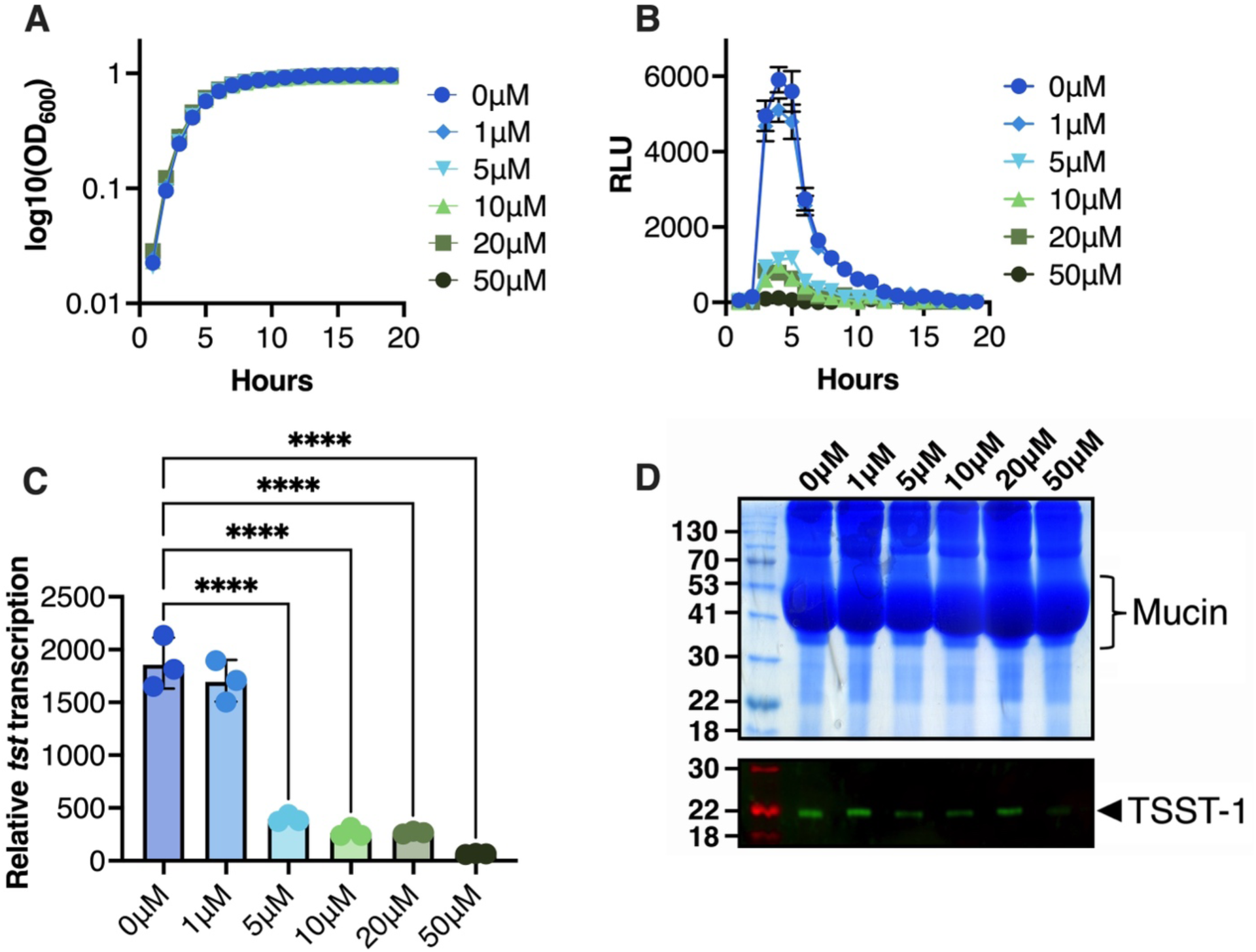
PP-HCl decreases *tst* promoter activity and TSST-1 production. (A) Growth of wild-type *S. aureus* MN8 containing pAmilux::P*_tst_* was assessed by optical density at 600 nm over 18 hours with parallel assessment of luminescence. Growth with various concentration of PP-HCl up to 50 µM was similar to untreated *S. aureus*. Results are presented as log10 of the optical density at 600 nm. (B) Luminescence from wild-type *S. aureus* MN8 containing pAmilux::P*_tst_* (measured in RLU) decreased drastically by 5 μM of PP-HCl and was almost absent at 50 μM PP-HCl. (C) Relative expression of *tst* promoter was calculated as the area under the curve of luminescence over the area under the curve of OD_600_ and demonstrates the same tendency as shown in the luminescence curves. The experiment was repeated with 3 biological replicates and error bars represent SEM. Ordinary one-way ANOVA was performed (****, *p* < 0.0001). (D) TSST-1 production in the supernatants of wild-type *S. aureus* MN8 was evaluated by Western Blot at the same concentrations of PP-HCl as tested during the luciferase assay. Supernatants were concentrated using trichloroacetic acid and normalized for all samples to reach 12 OD_600_ units. Shown are exoprotein profiles (top panels) and Western blot analysis (bottom panels) of TSST-1 for wild-type *S. aureus* MN8.

**Table 1.** Bacterial strains used in this study.

| Strain | Description | Source or reference |
| --- | --- | --- |
| ***S. aureus*** |  |  |
| MN8 | Prototypic menstrual TSS strain, *tst*+ | Blomster-Hautamaa et al, 1986 |
| MN8Δ*ccpA* | MN8 with deletion of *ccpA* gene | Dufresne et al, 2022 |
| MN8Δ*rot* | MN8 with deletion of *rot* gene | Tuffs et al, 2019 |
| MN8Δ*agr* | MN8 with the *agr* operon replaced with a *tetR* marker | Li et al, 2011 |
| MN8Δ*srrAB* | MN8 with deletion of *srrAB* genes | Tiwari et al, 2020 |
| MN8Δ*sarA* | MN8 with deletion of *sarA* gene | Baroja, 2016 |
| MN8Δ*saeS* | MN8 with deletion of *saeS* gene | Baroja, 2016 |
| MN8 (pAmilux::*Ptst*) | MN8 containing pAmilux::*Ptst* | Li et al, 2011 |
| MN8Δ*ccpA* (pAmilux::*Ptst*) | MN8Δ*ccpA* containing pAmilux::*Ptst* | Dufresne et al, 2022 |
| MN8Δ*rot* (pAmilux::*Ptst*) | MN8Δ*rot* containing pAmilux::*Ptst* | Tuffs et al, 2019 |
| MN8Δ*agr* (pAmilux::*Ptst*) | MN8Δ*agr* containing pAmilux::*Ptst* | Tuffs et al, 2019 |
| MN8Δ*srrAB* (pAmilux::*Ptst*) | MN8Δ*srrAB* containing pAmilux::*Ptst* | Tiwari et al, 2020 |
| MN8Δ*sarA* (pAmilux::*Ptst*) | MN8Δ*sarA* containing pAmilux::*Ptst* | Baroja, 2016 |
| MN8Δ*saeS* (pAmilux::*Ptst*) | MN8Δ*saeS* containing pAmilux::*Ptst* | Baroja, 2016 |
| ***Lactobacillus* species** |  |  |
| *Lactobacillus crispatus* ATCC 33820 | Representative strain of community state type I (CST I) | ATCC |
| *Lactobacillus gasseri* ATCC 33323 | Representative strain of CST II | ATCC |
| *Lactobacillus jensenii* ATCC 25258 | Representative strain of CST V | ATCC |

### Effect of PP-HCl on vaginal lactobacilli and T cell activation

The vaginal environment is considered to be strongly influenced by a beneficial microbiota (22), and the vaginal microbiome in women is generally classified into five Community State Types (CSTs). CST-I, CST-II and CST-V, dominated by *Lactobacillus crispatus*, *Lactobacillus gasseri* and *Lactobacillus jensenii*, respectively, are considered to provide a protective function (36); thus, an important property of PP-HCl for use as an anti-virulent therapeutic would be the lack of activity against beneficial microbiota members that may also play an important role in preventing mTSS (37). To evaluate this, we assessed growth of 3 representative beneficial vaginal *Lactobacillus* species in VDM containing increasing concentrations of PP-HCl (**Fig.3A-C**). Bacterial growth was similar in the presence of PP-HCl, and although there was a small growth defect with *L. crispatus* when grown in the presence of 250 µM PP-HCl, the compound did not inhibit growth of either *L. gasseri* or *L. jensenii*, further indicating that PP-HCl does not possess overt antibiotic properties against Gram positive bacteria (**Fig.3A-C**). As acidification of the vaginal environment is considered a key property of healthy CSTs (22), we assessed production of lactic acid in the presence of PP-HCl from the three lactobacilli species. Lactic acid production was similar for both treated and untreated conditions demonstrating that representatives of a healthy vaginal microbiota are not drastically affected by the presence of the anti-virulent compound (**Fig.3D**).

**Figure 3.**
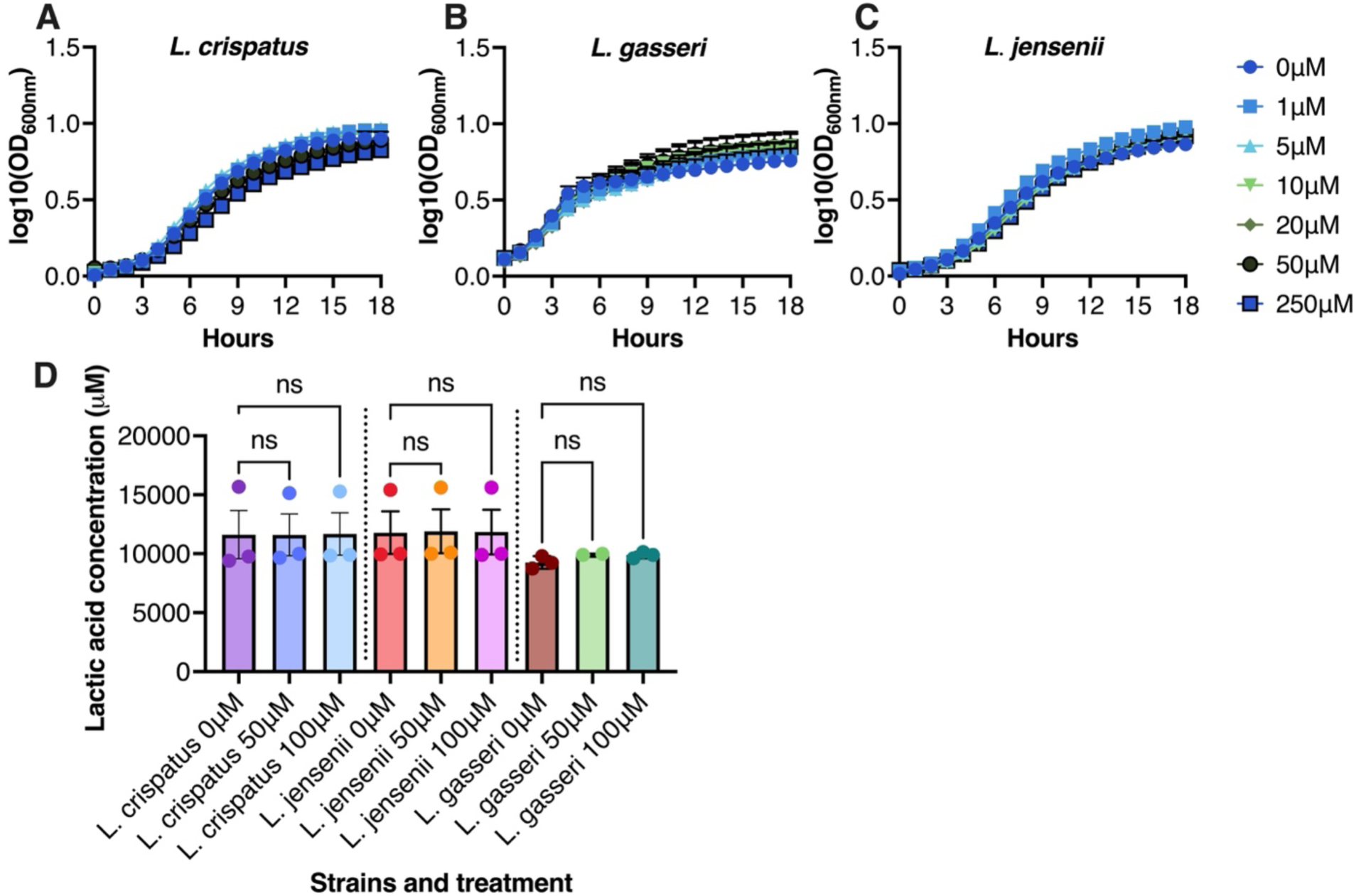
PP-HCl does not disrupt representative healthy members the vaginal microbiota. Growth and lactic acid production from the three dominant representativesof healthy or stable microbiota communities were assessed. (A) *L. crispatus* (ATCC 33820), (B) *L. gasseri* (ATCC 33323) and (C) *L. jensenii* (ATCC 25258) growth was assessed in 60mM glucose VDM at concentrations of PP-HCl ranging from 0 to 250 μM. (D) Lactic acid production by the lactobacilli at 0, 50 and 100 μM of PP-HCl was assessed and no differences were detectable within the groups. Ordinary one-way ANOVA was performed and *p* value over 0.05 were considered non-significant (ns).

Given that PP-HCl is used as an analgesic for urinary tract infections (38), we next evaluated PP-HCl for activity on eukaryotic cells with a focus on immune cells. To test this, human peripheral blood mononuclear cells (PBMCs) were isolated from healthy donor blood and treated with either purified recombinant TSST-1, PP-HCl, or in combination. As expected, TSST-1 induced production of the T cell cytokine IL-2 whereas PP-HCl did not; however, in combination the IL-2 response was reduced relative to TSST-1 alone (**Fig. S1A**). We also assessed cell viability and TSST-1 treatment showed decreased cell viability, likely due to activation induced cell death, whereas PP-HCl did not affect cell viability. In combination, cell viability was also decreased similar to TSST-1 alone (**Fig. S1B)**. We then assessed supernatants from wild-type *S. aureus* MN8, or the *S. aureus* MN8 with a deletion in the TCS SaeRS. SaeRS is a direct and positive regulator of the *tst* promoter and an in-frame deletion in the histidine kinase gene *saeS* results in a drastic reduction of TSST-1 production (35). Wild-type *S. aureus* MN8 supernatant induced the production of IL-2 which was significantly reduced when using the Δ*saeS* mutant and was similar to co-treatment of PBMCs with the wild-type *S. aureus* MN8 supernatant and PP-HCl (**Fig. S1C**). None of the supernatants, or PP-HCl alone, altered cell viability in between the various conditions tested (**Fig. S1D**). These experiments demonstrate that in addition to the ability of PP-HCl to reduce TSST-1 expression from *S. aureus*, this compound also appears to inhibit superantigen-induced T cell activation to levels reached by the *S. aureus* MN8 Δ*saeS* mutant.

### Activity of PP-HCl on TSST-1 transcription bypasses the main tst repressors

To decipher genetic mechanisms involved in PP-HCl-dependent repression of TSST-1, we evaluated various key *S. aureus* MN8 regulatory mutant strains containing pAmilux::P*_tst_* in the presence or absence of PP-HCl. As before, 50µM PP-HCl had little impact on *S. aureus* growth (**Fig.4A and 4C)**. The two known positive regulators of *tst* transcription are the Agr and Sae TCSs where SaeR acts directly on the *tst* promoter, while Agr relieves repression of *tst* by Rot (24, 35). As predicted, *tst* expression was dramatically reduced in the *agr* and *saeRS* mutants without PP-HCl treatment (**Fig.4B and 4E**). Key repressor systems for *tst* transcription in rich media include the two component system SrrAB (9, 39), and the cytoplasmic intermediatory regulators *ccpA*, *rot*, and *sarA* (12, 24, 35); however, in the tested conditions, only the Δ*ccpA* mutation showed an increase in *tst* expression (**Fig.4E**). Surprisingly, the other known repressors (SrrAB, Rot and SarA), in VDM with low levels of glucose, had decreased *tst* expression suggesting that may fulfill an activator role in this specific environment instead, and that other levels of regulation are yet to be described in these mTSS conditions (**Fig.4E**). PP-HCl completely inhibited luminescence within the various strains suggesting that it repressed *tst* expression in each of these mutant backgrounds (**Fig.4D**). The lack of TSST-1 transcription from these latter regulators in the presence of PP-HCl suggests that the compound does not function by activating these repressors (**Fig.4E**).

**Figure 4.**
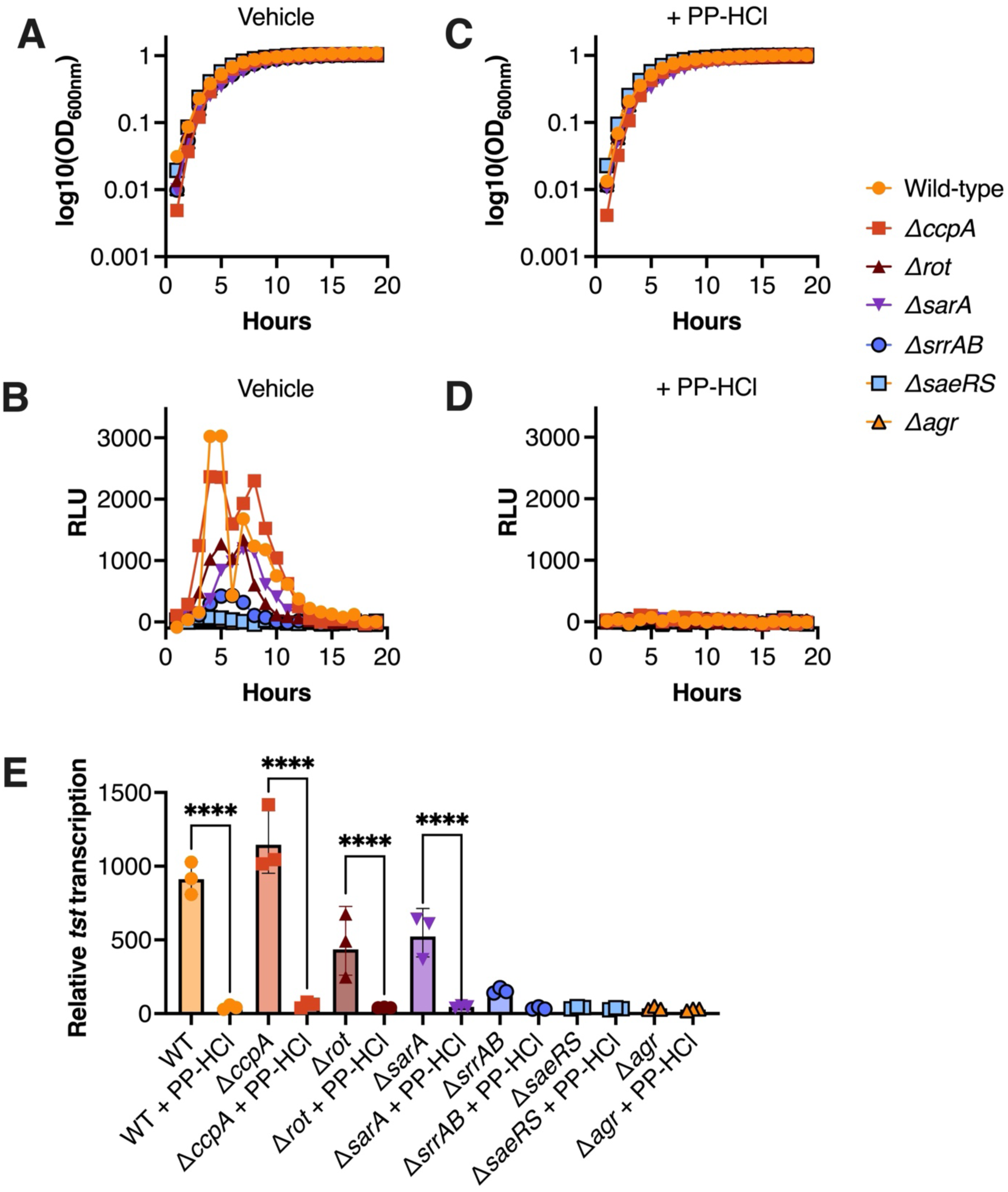
PP-HCl overcomes known genetic regulators of TSST-1 to inhibit transcription of *tst*. Growth curves and luciferase assays of *S. aureus* MN8 reporter strains including central TSST-1 regulatory mutants was performed (A, B) without or (C, D) with 50 μM of PP-HCl. (E) The relative expression of the *tst* promoter in each mutant exposed to both conditions are presented. The experiment was repeated with 3 biological replicates. Ordinary one-way ANOVA was performed for statistical analysis (****, *p* < 0.0001).

### PP-HCl-dependent inhibition of multiple exotoxin genes requires the SaeRS TCS

To further evaluate potential regulatory circuits involved in the PP-HCl-mediated repression of TSST-1, we conducted transcriptional analyses by RNA sequencing comparing wild-type *S. aureus* MN8 grown for 4h in the presence or absence of 50 µM PP-HCl (**Fig.5A**). This analysis demonstrated that the TSST-1 gene, as well as several exotoxin genes, were repressed in the presence of PP-HCl (**Fig.5A, Table S1**). Interestingly, α-toxin, TSST-1, Sbi, SCIN and CHIPs are all known to be positively regulated by the SaeRS TCS (35, 40) and each of these genes, including the gene for the more recently charactered myeloperoxidase inhibitor SPIN (41), all contain canonical SaeR binding sequences upstream of the respective promoters (42). Although the α-hemolysin toxin gene in *S. aureus* MN8 is truncated (denoted as α-hemolysin*), the transcript is produced and was highly repressed in the presence of PP-HCl. Although not statistically different, we also noted a trend for decreased transcription of genes encoding the gamma-hemolysin (*hlgA*, *hlgB*, *hlgC*), also known to be SaeRS controlled (42), as well as both *saeR* and *saeS* (**Fig. 5A and C**), the latter changes are likely due to the known autoregulation of the *sae* locus. Of genes that could be involved in virulence, we also noted the upregulation of *sdrE*. These data suggested that repression of *tst* transcription by PP-HCl, and the noted exotoxins, maybe be mediated through inhibition of the SaeRS TCS.

**Figure 5.**
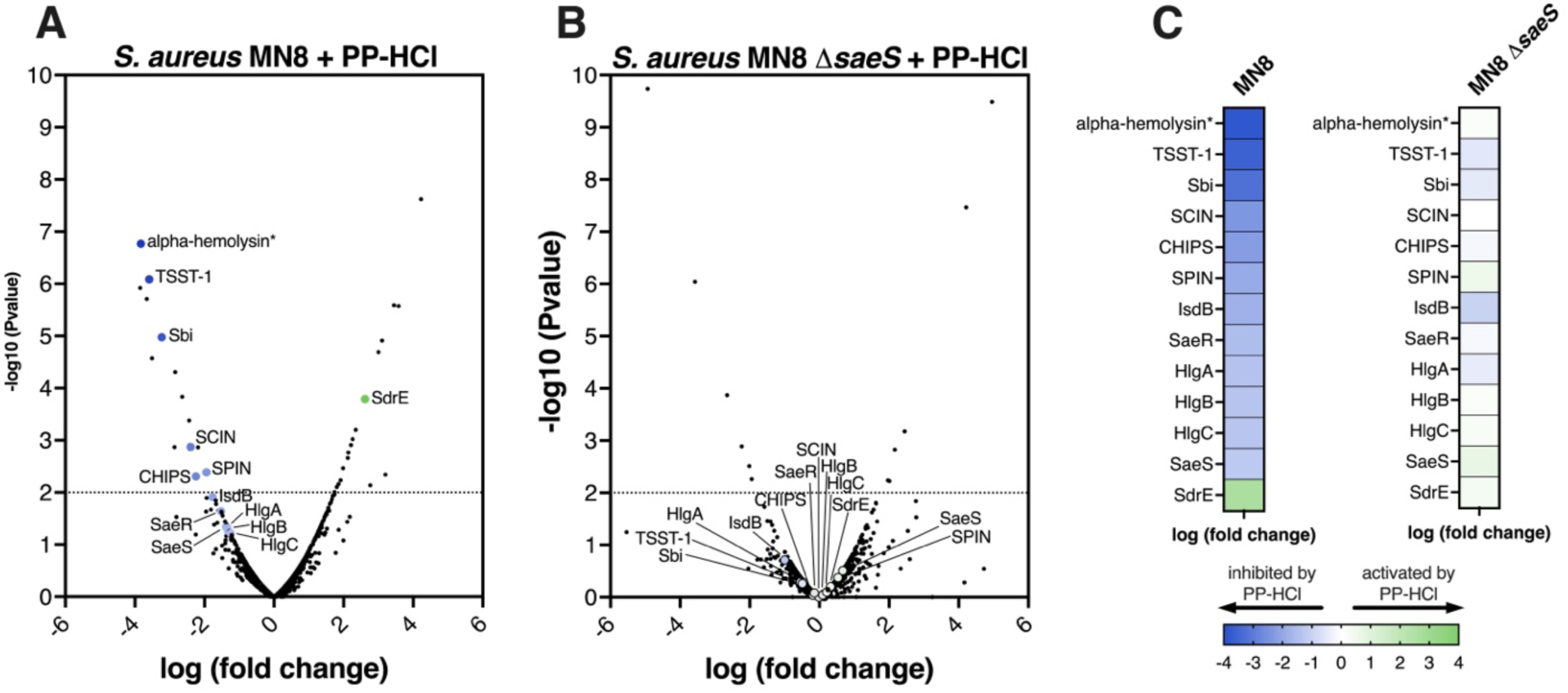
PP-HCl affects transcriptional regulation of a variety of virulence factors related with SaeRS regulation. RNAseq comparing *S. aureus* MN8 untreated versus MN8 treated with 50 μM of PP-HCl. See Table S1 for the extensive analysis of transcripts (A). The same analysis was repeated for MN8 Δ*saeS* in absence and presence of PP-HCl. See Table S2 for the extensive analysis of transcripts (B). TSST-1 and several other virulence factors are either inhibited or activated in presence of the compound in the wild-type strain: these various factors are represented in a scale of blue, if inhibited, to green when activated. The list of these significant changes associated with virulence are presented for the wild-type strain and compared to the transcriptional changes of the same genes in MN8 Δ*saeS* (C).

To evaluate the involvement of SaeRS in the inhibition of virulence factor expression by PP-HCl, the RNAseq experiment was repeated in the *S. aureus* MN8 Δ*saeS* mutant in the presence or absence of PP-HCl (**Fig.5B**). Transcription of the genes encoding each of the exotoxins was not differentially changed in the Δ*saeS* genetic background between the two conditions (**Fig. 5B, Table S2**). These data indicate the importance of SaeRS pathway for PP-HCl to repress TSST-1 and the other exotoxins (**Fig. 5C**).

### PP-HCl inhibits phosphorylation of SaeS

SaeS is the sensor histidine kinase of the SaeRS TCS that relays the presence of polymorphonuclear leukocytes (PMN or neutrophil)-produced signals via phosphorylation to its response regulator SaeR, which subsequently binds target DNA to alter gene transcription (40, 43, 44). Preynat-Seauve *et al* proposed that PP-HCl could inhibit human kinases by binding the ATP-binding site (45) and therefore we hypothesized that PP-HCl may bind the ATP-binding site of SaeS to inhibit phosphorylation of SaeR. To evaluate this, *S. aureus* MN8 was grown in the presence or absence of PP-HCl and the phosphorylated state of SaeR was assessed (**Fig. 6A**). Relative to untreated wildtype *S. aureus* MN8 cells, PP-HCl inhibited phosphorylation of SaeR that was similar to the MN8 Δ*saeS* deletion mutant, that was more prominent by 18h (**Fig. 6A**). Using Sortase A (SrtA) as an internal normalization control, PP-HCl decreased the expression of SaeR at both 4 and 18 hours of incubation that was similar to the MN8 Δ*saeS* strain (**Fig. 6B & 6C**). PP-HCl also reduced phosphorylation of SaeR compared with untreated wildtype cells, whereas phosphorylated SaeR was not detectable in the MN8 Δ*saeS* deletion mutant (**Fig. 6D & 6E**). As these results indirectly suggested that PP-HCl may potentially interfere with the ATP-binding site of SaeS, a competition assay in between ATP and PP-HCl in presence of purified SaeS and SaeR was designed. Low levels of ATP were not sufficient to compete against PP-HCl (**Fig. 6F, compare lanes 4 and 5**) whereas increased ATP concentrations was able to counteract the inhibition of PP-HCl (**Fig. 6F, compare lanes 5 and 7)**. Taken together, these data demonstrate that PP-HCl can inhibit phosphorylation of SaeR and further suggests that PP-HCl may compete for the ATP-binding site of SaeS.

**Figure 6.**
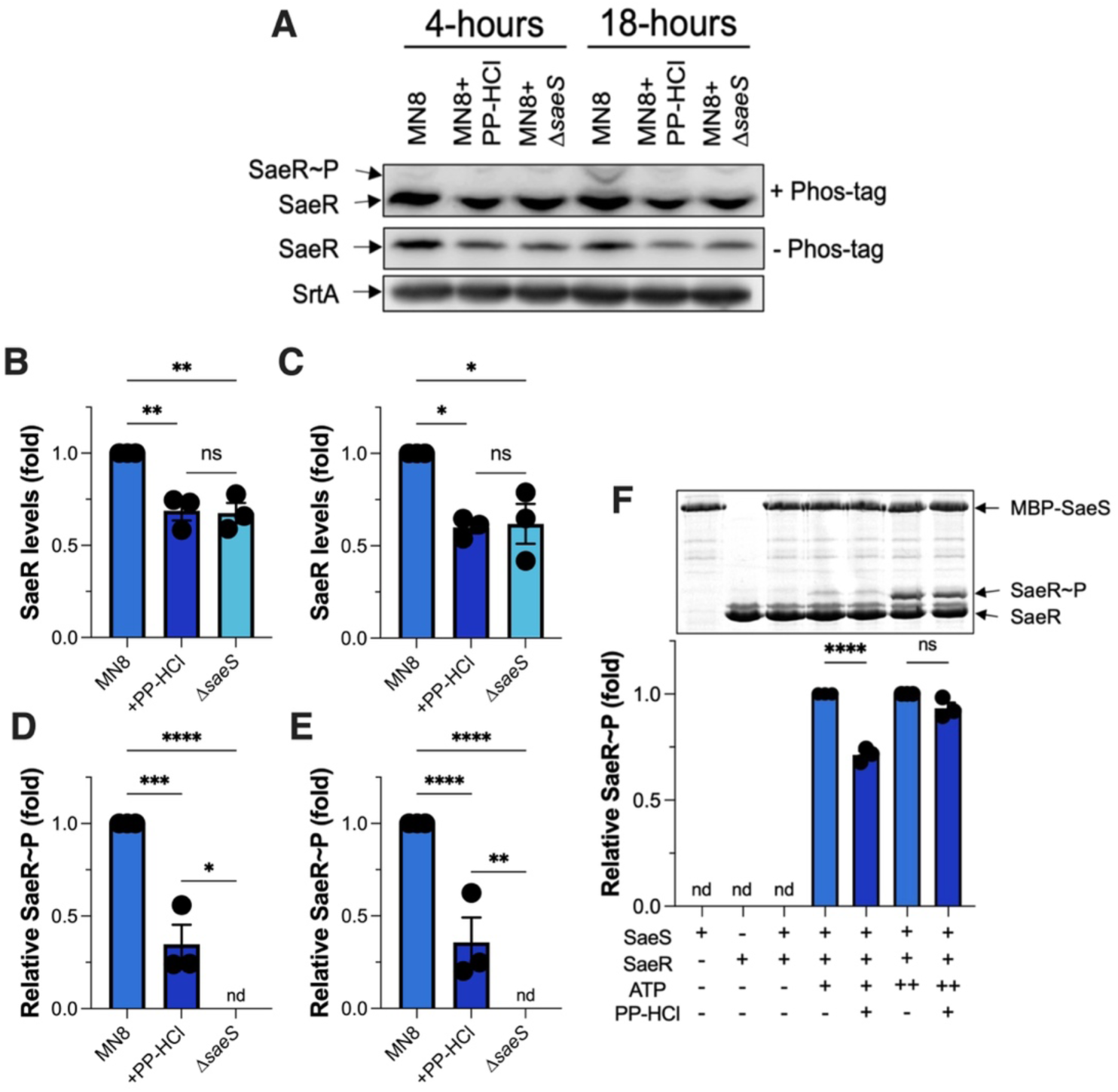
PP-HCl targets the SaeRS TCS through competition for the ATP-binding site of SaeS. Phosphorylated SaeR levels were assessed using Phos-tag Western Blot both at 4- or 18-hours incubation of *S. aureus* MN8 strains in the various conditions. *S. aureus* MN8 Δ*saeS* was used as negative control for phosphorylation of SaeR. (A) Representative figure of the biologically triplicated experiment is presented. (B-C) Calculations of relative proteinaceous levels are analyzed against sortase A (SrtA). Total SaeR levels are compared at 4- and 18-hours incubation respectively. Ordinary one-way ANOVA was performed (ns, *p* > 0.05; *, *p* ≤ 0.05; **, *p* value < 0.01). (D-E) The relative phosphorylated SaeR levels in between untreated and treated *S. aureus* MN8 are compared at the same time points. Ordinary one-way ANOVA was performed (****, *p* < 0.0001). (F) Phosphorylation assay of recombinant SaeR by MBP-SaeS in competition of PP-HCl with ATP tested in near Km ATP concentration (200 μM; indicated by +) or in excess ATP (10 mM; indicated by ++). Ordinary one-way ANOVA was performed.

## Discussion

Prevention of mTSS has been a concern since the epidemic from the early 1980s (11), and although the incidence of mTSS has since remained relatively low, outcomes from mTSS cases can be devastating and life-threatening (13). Preventing the production of TSST-1 during menses is important to avoid mTSS and product components must be evaluated to promote tampon safety, including their biocompatibility and chemical safety, impact on the vaginal mucosa and microbiota, as well as effects on *S. aureus* growth and TSST-1 production (46). Nevertheless, multiple conditions can still occur during the menstrual cycle allowing for the overproduction of TSST-1 to result in mTSS. In order to expand approaches to further prevent mTSS, we developed a screen to discover anti-virulence molecules that inhibit TSST-1 production from *S. aureus* (**Fig. 1**). This screen used the prototypical mTSS strain *S. aureus* MN8 (21) coupled with growth in an environment designed to mimic vaginal conditions that could lead to mTSS (12). Here, we report the discovery of PP-HCl as a potential new anti-virulence molecule that represses exotoxin production in *S. aureus* by inhibiting kinase activity of the SaeRS TCS.

In this study, we mimicked the conditions of the vaginal tract during mTSS including increased oxygenation, near neutral pH, and decreased glucose content. In these conditions, we noted differential activity of known regulators of the toxin TSST-1. Compared to prior experiments where the media were not adapted to the environmental cues related with mTSS, mutations in Rot, SarA and SrrAB reduced overall *tst* expression whereas in standard laboratory media these function as repressors **(Fig. 4)** (20, 24, 35). These data solidify our previous findings that CcpA is the main repressor within the vaginal niche and that decreasing glucose levels during menses explains the de-repression of TSST-1 within this short window (12). This also highlights the intricate nature of regulation networks within a bacterium and the importance of mimicking the disease-prone environment when studying the expression of virulence factors.

PP-HCl is an azo dye and analgesic that is used for the treatment of symptomatic urinary tract infections (38) and this compound was associated with inhibition of phosphatidylinositol kinases involved in nociception, explaining its analgesic effects (45). The cytoplasmic component of bacterial histidine kinases, including SaeS, generally contain 2 conserved domains: one domain is involved in both autophosphorylation and phospho-transfer to a cognate response regulator, whereas the second domain acts as the catalytic ATP-binding domain (47). Following the *S. aureus* RNAseq experiments in the presence of PP-HCl that suggested a role for SaeRS in the exotoxin repressive phenotype (**Fig. 5**), we found that phosphorylation of SaeS to its cognate response regulator SaeR is reduced in the presence of PP-HCl both in *S. aureus* (**Fig. 6A-E**) and using recombinant SaeS and SaeR proteins (**Fig. 6F**). Excess ATP was able to overcome inhibition of SaeS by PP-HCl suggesting the compound may interfere with ATP binding (**Fig. 6F**) although future studies are focused on understanding how PP-HCl inhibits this activity.

Although our screen targeted the inhibition of TSST-1 production and identified the SaeRS system as the major pathway targeted by PP-HCl, the transcriptional changes may not be exclusive to SaeRS-dependent gene regulation. From the RNAseq experiments **(Fig 5A and Tables S1-S3**) we focused on altered expression of known virulence factors, and although multiple exotoxins were repressed, we also noted increased expression in *sdrE* transcripts in the presence of PP-HCl (**Fig. 5A**) that were no longer induced in the MN8 Δ*saeS* strain (**Fig. 5B**). However, comparing wild-type *S. aureus* MN8 with the Δ*saeS* mutant in the absence of PP-HCl, *sdrE* transcripts were not markedly altered (**Fig. S2**), and furthermore, we did not observe a canonical SaeR binding motif (42) upstream of the *sdrE* gene. SdrE is a *S. aureus* surface protein that functions to inhibit complement activation by high-affinity binding of the complement regulatory protein Factor H (48), and thus PP-HCl could potentially enhance virulence under certain situations. Nevertheless, most PP-HCl repressed virulence factors overlapped with repression in the Δ*saeS* mutant in the absence of PP-HCl (**Fig. S2)**, and these repressed virulence factors were not altered in the Δ*saeS* mutant in the presence of PP-HCl (**Fig. 5**).

Anti-virulent molecules targeting sensing systems from *S. aureus* have already shown great potential. As previously mentioned, TCSs including the Agr system, ArlRS, GraRS and also SaeRS, have been subjects of prior investigation. In the context of preventing risks of mTSS, we demonstrated previously that repression of TSST-1 is mainly due to glucose sensed through CcpA (12) and, conversely, that TSST-1 is predominantly activated by SaeRS (35). We consider SaeRS to be an exceptional target as its activation has been associated with the positive regulation of several adhesins and toxins that results in attenuated virulence *in vivo* (35, 49–53). Furthermore, the IsdB iron uptake system was downregulated in the presence of PP-HCl and this system is important for colonization of *S. aureus* USA300 in the mouse vaginal tract (54). Another advantage for targeting SaeRS compared to the Agr TCS is that no attenuated or mutated SaeRS strain has been isolated in clinics to our knowledge (55). We hypothesize that such strains would become avirulent and less prone to colonize the vaginal niche due to a modification of the SaeS ATP-binding site that would lead to reduced SaeR phosphorylation and subsequently reduced toxin and adhesin expression. Altogether, SaeRS may be a primary choice for targeting superantigen-specific diseases such as mTSS. A key advantage of targeting virulence rather than bacterial growth is to limit antibacterial off-target activity which may otherwise lead to expansion of antimicrobial resistance (56). Indeed, PP-HCl did not induce killing of *S. aureus* (**Fig. 2**), or key representative species of vaginal lactobacilli (**Fig. 3**) that may also be advantageous for preventing mTSS (37). Thus, PP-HCl may represent a new lead compound and strategy to further develop safer menstrual products to prevent the occurrence of mTSS.

## Experimental procedures

### Ethics Statement

Human blood from healthy donors was obtained in accordance with the human subject protocol HSREB 110859 approved by the London Health sciences centre (LHSC) research ethic board at the University of Western Ontario. Volunteers were recruited by passive advertising through the Department of Microbiology and Immunology at the University of Western Ontario and all volunteers gave a written informed consent before each sampling. Each sample is fully anonymized and no information regarding the identity of the donor was retained.

### Bacterial strains and high-throughput screening of bioactive molecules

The list of strains used for this study is found in Table 1. Routine growth of *S. aureus* MN8 and derivatives was done aerobically at 37°C in tryptic soy broth (TSB) with shaking at 250 rpm supplemented with the appropriate antibiotics as needed. For experiments measuring the production of TSST-1, *S. aureus* strains were grown in vaginally defined medium (VDM) (57) modified to contain 700 μM glucose to mimic conditions favorable for the production of TSST-1 (12). The luciferase screen used *S. aureus* MN8 harbouring P*_tst_* reporter in fusion with the *lux* operon (pAmilux::P*_tst_*). Briefly, bacteria were grown and the OD_600_ was adjusted to 0.01 in fresh media and distributed in 384-wells microplates using the Tempest dispenser (Formulatrix). Expression control was the reporter strain in media of assay and the repression control was the reporter strain in assay media containing 60mM glucose known to repress TSST-1 (12). Small molecules were pre-inoculated (at 10μM) before adding the bacterial inoculum and 3920 molecules were tested. Both luminescence and OD_600_ were measured every hour for 18 hours during incubation at 37°C with continuous agitation. For each molecule, the point with the highest luminescence was compiled with its respective OD_600_. Seventy initial compounds with normalized growth equal to or better than 95% of the expression control and with raw luminescence lower than 100 RLU were selected. Known antimicrobials in this list were removed as the goal of the study is to identify molecules with anti-virulent but not antimicrobial properties.

*S. aureus* MN8 containing pAmilux::P*_tst_* were grown as above and subsequently challenged with remaining compounds and titrated from 0μM to 50μM and both luminescence and OD_600_ were measured every hour for 18 hours in a Biotek Synergy H4 multimode plate reader. Relative luminescence units (RLUs) were calculated from the area under the curve of the luminescence over the area under the curve of the absorbance during the same time. For later assays, only 0 and 50μM PP-HCl in 700 μM glucose VDM were used with the various *S. aureus* MN8 strains. Western immunoblots for TSST-1 production were performed as previously described (35).

### RNA sequencing experiments

RNAseq was performed as described previously (12, 19). Briefly, *S. aureus* MN8 or MN8 Δ*saeS* were grown with 0 or 50μM PP-HCl in 700 μM glucose VDM for 4 hours, RNA was extracted using RNeasy Plus Mini kit (QIAGEN) and subsequently treated with Turbo DNA-free Kit (Ambion). RNA sequencing and its comparative analysis was performed by SeqCenter (Pittsburgh, USA). Twelve million paired-end Illumina sequencing was performed and followed by analysis as previously described (12). Raw RNA data were deposited at NCBI under BioProject accession no. PRJNA1080564.

### CST growth and lactic acid production

Growth of lactobacilli was assessed as previously described (19). All representatives of stable/healthy Community State Types (CST) were grown overnight in Man, Rogosa and Sharpe (MRS) media, then subcultured in 60mM glucose VDM supplemented with or without 50μM PP-HCl at a starting OD_600_ of 0.05. Cultures were incubated for 20 hours at 37°C without agitation. Growth was assessed by OD_600_ readings every hour in a Biotek Synergy H4 multimode plate reader. Lactic acid production was assessed for each representative strain after 24 hours incubation in the same conditions as for the growth assessment using the Lactate-Glo Assay (Promega) following the manufacturer’s protocol.

### IL-2 quantification and viability of PBMC

Peripheral blood mononuclear cells (PBMC) were isolated with Ficoll Paque Plus following the manufacturer protocol and seeded for a final concentration of 1 x 10^6^ cells/ml in 96-well plates. Cells were either challenged for 18 hours at 37°C and 5% CO_2_ against *S. aureus* MN8 or MN8 Δ*saeS* filtered supernatants exposed to 0 and 50μM PP-HCl, or with 100 ng/ml of purified recombinant TSST-1 (24). IL-2 was measured by human IL-2 ELISA (Invitrogen) using the manufacturer instructions. Data at a dilution factor of 6250 were normalized over the control condition and plotted in biological triplicates. Results are presented as the percentage of IL-2 production compared to their respective controls.

Viability of PBMCs was assessed by incubating the challenged cells as above an additional hour with the resazurin-based PrestoBlue HS Cell Viability Reagent (Invitrogen) following the manufacturer instructions and plates were read for fluorescence at excitation of 560 nm and emission of 590nm and for absorbance of 570 nm in biological triplicates. Results are presented as the percentage viability compared to their respective controls.

### Determination of SaeR phosphorylation state

*S. aureus* MN8 was grown in 700 μM glucose VDM supplemented with or without 50μM PP-HCl and incubated for 4 or 18 hours at 37°C with agitation. *S. aureus* MN8 Δ*saeS* was grown in the same conditions as a control for SaeR phosphorylation. Separation of SaeR and SaeR∼P was performed as described previously using 12% polyacrylamide gels containing the acrylamide-pendant Phos-tag ligand (50). Briefly, whole cell extracts were obtained by resuspending cell pellets in cell extract buffer (20 mM Tris [pH 7.0], 1× Protease Inhibitor Cocktail Set I (Sigma-Aldrich)) and transferred to sterile screw cap tubes containing silica beads. Cells were homogenized at room temperature using a Precellys 24 bead beater (Bertin technologies) for 3 cycles of 6500 m/s, 30 s each, and centrifuged. Whole-cell extracts were normalized by protein concentration (A_280_) to 100 µg and electrophoresed on Phos-tag gels with standard running buffer (0.1% [w/v] SDS, 25 mM Tris, 192 mM glycine) at 4°C under constant voltage (150 V) for 2 h. Gels were washed for 15 minutes with transfer buffer (25 mM Tris [pH 8.3], 192 mM glycine, 20% methanol) containing 1mM EDTA followed by a second wash without EDTA to remove manganese ions. Proteins were then transferred to PVDF membranes (Cytiva) and incubated with polyclonal rabbit antibodies to SaeR (1.5:1,000) for 1 h. Membranes were then washed with TBST and incubated with StarBright Blue 700 goat anti-rabbit IgG (1:3500; Bio-Rad) for 1 h. Membranes were washed in TBST and signals were visualized using an Amersham ImageQuant800. The densities of the SaeR∼P signal were determined by quantification with Multi Gauge software (FujiFilm). The data are representative of three independent experiments, and a representative image is shown.

*In vitro* kinase assays using recombinant protein were performed as previously described (50). To test the effect of PP-HCl on kinase activity *in vitro*, 50µM PP-HCl was added to the standard reaction mixture (5 µM MBP-SaeS, 10 µM SaeR-His_6_, 200 µM ATP, 1× TKM buffer) and incubated at 37°C for 1 h. the reaction was stopped by addition of 5× SDS loading buffer. Phosphorylated and unphosphorylated forms of SaeR were separated using 12% phos-tag acrylamide gels and visualized by Coomassie blue staining. The resulting gels were imaged using an Amersham ImageQuant800, and the levels of SaeR∼P were determined by quantification with Multi Gauge software.

### Statistical analysis

Statistical analysis was performed using GraphPad Prism 10. Ordinary one-way ANOVA was used without correction for multiple comparisons unless mentioned otherwise.

## Acknowledgements

This work was supported by funding from the Canadian Institutes of Health Research (CIHR) Grant PJT-166050 to JKM and by funding from the National Institutes of Health (NIH) Grant R01 AI137403 to SRB. We acknowledge Dr. Eric Brown and the Centre for Microbial Chemical Biology (CMCB) at McMaster University, including Tracey Campbell, Susan McCusker, and Cecilia Murphy, for their support and advice with the screening of bioactive molecules.

---

**Figure S1.**
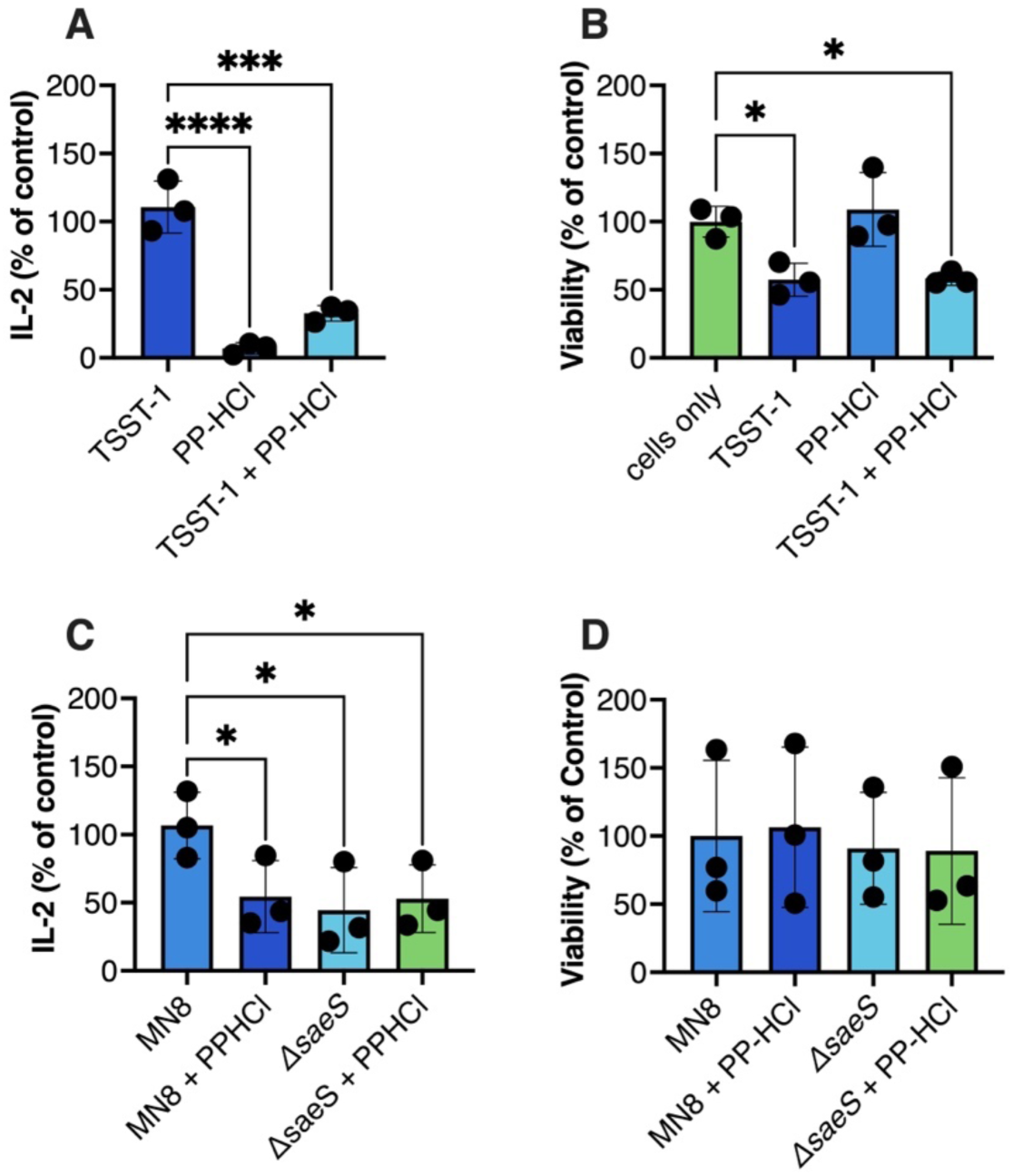
PP-HCl decreases TSST-1-induced IL-2 production from PBMC. (A) Percentage of IL-2 compared to control condition (100 ng/ml TSST-1) was measured for each condition. Ordinary one-way ANOVA was performed (***, *p* < 0.001; ****. *p* < 0.0001). (B) Percentage of viability compared to the control condition (cells only) was measured for each condition. Ordinary one-way ANOVA was performed (*, *p* ≤ 0.05). (C) Percentage of IL-2 production at a dilution factor of 6250 was compared to the *S. aureus* MN8 supernatant control. Ordinary one-way ANOVA was performed. (D) Percentage of viability compared to the *S. aureus* MN8 supernatant condition was measured for each condition. Ordinary one-way ANOVA was performed (D).

**Figure S2.**
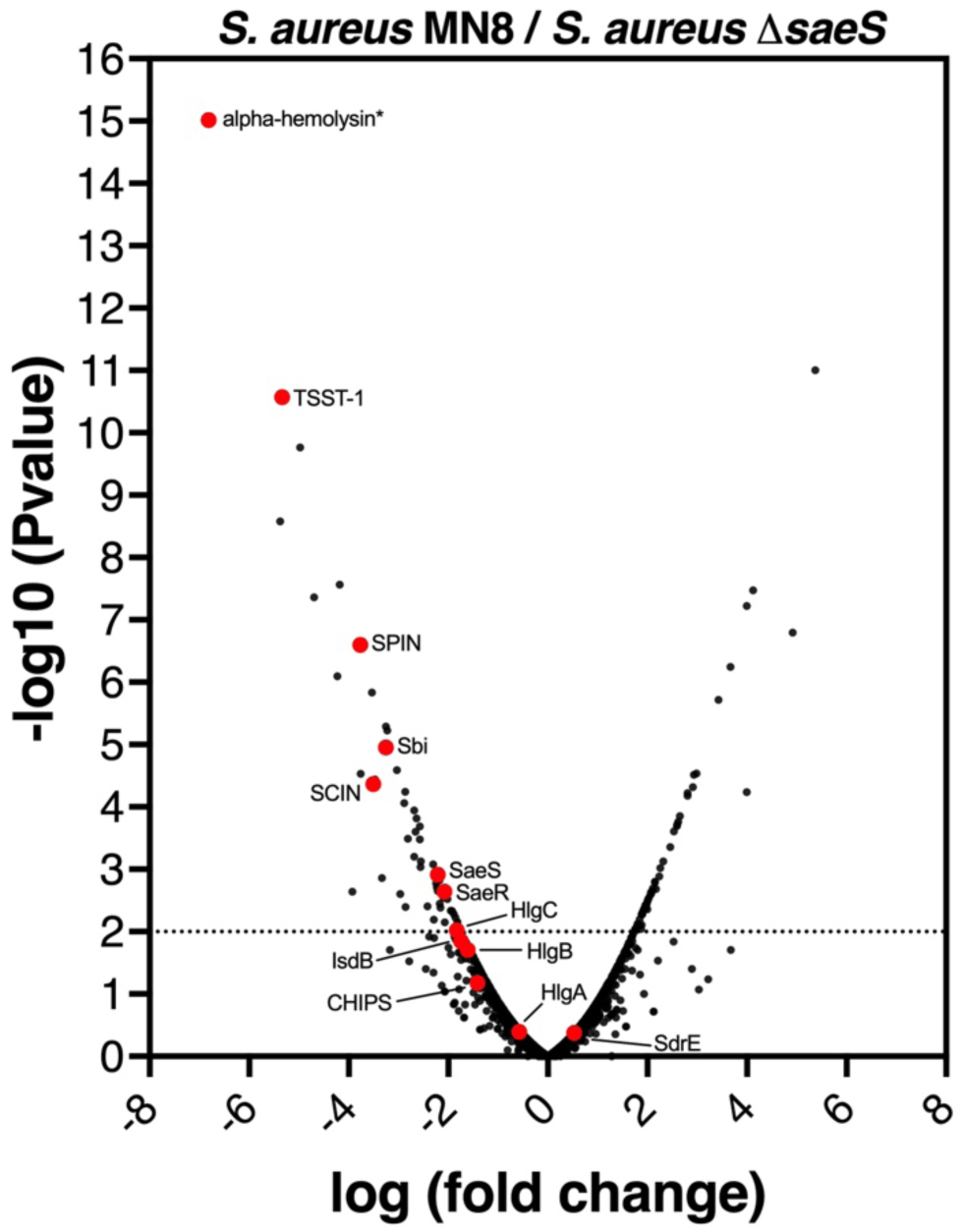
Inactivation of SaeRS regulon decreases virulence of *S. aureus* MN8 similar to addition of PP-HCl. RNAseq comparing *S. aureus* MN8 versus MN8 Δ*saeS* in 700 μM glucose VDM was analyzed. The main virulence factors associated with SaeRS regulon, also identified in Figure 5, are presented with red dots. See Table S3 for the extensive analysis of transcripts.

**Table S1..** Significant transcriptional comparison analysis of all differentially expressed genes in between MN8 with or without PP-HCl.

| Locus tag | Gene name | Description | (log) Fold Change | P-value | False Discovery Rate |
| --- | --- | --- | --- | --- | --- |
| NLA26_RS00140 |  | chemotaxis-inhibiting protein CHIPS | -2,24302619 | 0,004935101 | 0,462313203 |
| NLA26_RS00160 |  | hypothetical protein | -3,659580928 | 1,95E-06 | 0,000852383 |
| NLA26_RS02710 | isdB | heme uptake protein IsdB | -1,769367438 | 0,012306881 | 0,896693043 |
| NLA26_RS02845 |  | fibrinogen-binding protein | -1,599693106 | 0,02215777 | 1 |
| NLA26_RS02870 | scc | complement inhibitor SCIN-C | -2,39882216 | 0,001344988 | 0,162433602 |
| NLA26_RS02885 | hyl | alpha-hemolysin | -3,828206866 | 1,70E-07 | 0,000149017 |
| NLA26_RS03150 |  | hypothetical protein | 1,740261214 | 0,00979148 | 0,778274301 |
| NLA26_RS04210 |  | phosphate ABC transporter substrate-binding protein PstS | 1,372391329 | 0,047233419 | 1 |
| NLA26_RS04785 |  | tail protein | 2,766440303 | 0,007289357 | 0,616773681 |
| NLA26_RS06340 |  | AAA family ATPase | -3,847933922 | 1,20E-06 | 0,000629896 |
| NLA26_RS06345 |  | DDE-type integrase/transposase/recombinase | 3,57819408 | 2,67E-06 | 0,000876259 |
| NLA26_RS06515 |  | IS3 family transposase | -2,859483473 | 0,001347849 | 0,162433602 |
| NLA26_RS07110 |  | DM13 domain-containing protein | -2,642615584 | 0,00014624 | 0,027399133 |
| NLA26_RS07115 |  | DoxX family protein | -2,439377481 | 0,000417208 | 0,068396028 |
| NLA26_RS07120 | saeR | response regulator transcription factor SaeR | -1,521562381 | 0,022923182 | 1 |
| NLA26_RS07285 |  | transposase | -2,806802315 | 0,029588432 | 1 |
| NLA26_RS07820 | adhP | alcohol dehydrogenase AdhP | 1,66546417 | 0,013414396 | 0,925946341 |
| NLA26_RS07845 |  | aldo/keto reductase | 3,001823827 | 2,04E-05 | 0,00486424 |
| NLA26_RS07985 |  | MFS transporter | 1,393914093 | 0,036328038 | 1 |
| NLA26_RS08035 | sdrE | MSCRAMM family adhesin SdrE | 2,616281758 | 0,000162171 | 0,028358307 |
| NLA26_RS08665 |  | ABC transporter permease | 2,173488812 | 0,029323086 | 1 |
| NLA26_RS08710 |  | DUF2294 domain-containing protein | 1,370404433 | 0,04093036 | 1 |
| NLA26_RS08715 |  | YbcC family protein | 2,347450254 | 0,000631615 | 0,097454556 |
| NLA26_RS08720 |  | NADH dehydrogenase subunit 5 | 3,108270586 | 1,23E-05 | 0,003218903 |
| NLA26_RS08805 | spn | myeloperoxidase inhibitor SPIN | -1,942376963 | 0,004116629 | 0,415304568 |
| NLA26_RS08830 |  | hypothetical protein | 11,97357639 | 4,96E-16 | 1,30E-12 |
| NLA26_RS08845 |  | superantigen-like protein SSL7 | -1,811789693 | 0,021411536 | 1 |
| NLA26_RS09055 |  | terminase small subunit | -1,627938062 | 0,038020906 | 1 |
| NLA26_RS09330 |  | PTS ascorbate transporter subunit IIC | 1,986597389 | 0,003435188 | 0,360419915 |
| NLA26_RS09460 |  | formate/nitrite transporter family protein | 1,804605025 | 0,007836755 | 0,642368993 |
| NLA26_RS09780 |  | PTS transporter subunit EIIC | 1,506045462 | 0,026572905 | 1 |
| NLA26_RS09785 |  | L-lactate dehydrogenase | 1,69511943 | 0,011536168 | 0,864553393 |
| NLA26_RS09800 |  | hypothetical protein | 1,537702319 | 0,022714696 | 1 |
| NLA26_RS09805 |  | DUF488 domain-containing protein | 1,911986743 | 0,005807458 | 0,525274541 |
| NLA26_RS09880 | pflA | pyruvate formate-lyase-activating protein | 1,510539169 | 0,024662812 | 1 |
| NLA26_RS09885 | pflB | formate C-acetyltransferase | 2,261360621 | 0,000949328 | 0,138338252 |
| NLA26_RS09910 |  | isoprenylcysteine carboxyl methyltransferase family protein | -3,504918267 | 2,69E-05 | 0,005870903 |
| NLA26_RS10110 |  | ABC transporter ATP-binding protein | 3,199782966 | 0,004550577 | 0,44208016 |
| NLA26_RS10225 | adhE | bifunctional acetaldehyde-CoA/alcohol dehydrogenase | 2,117597635 | 0,002181478 | 0,238417346 |
| NLA26_RS11185 |  | glutathione peroxidase | -1,407029256 | 0,036103814 | 1 |
| NLA26_RS11200 |  | NAD(P)-binding protein | -1,476126369 | 0,027856441 | 1 |
| NLA26_RS11210 |  | CitMHS family transporter | -2,842911684 | 4,94E-05 | 0,009961835 |
| NLA26_RS11290 |  | hypothetical protein | 1,318450015 | 0,049944002 | 1 |
| NLA26_RS11645 |  | hypothetical protein | 1,587481609 | 0,022400495 | 1 |
| NLA26_RS11695 |  | D-lactate dehydrogenase | 1,558244497 | 0,019759696 | 1 |
| NLA26_RS11720 |  | ring-cleaving dioxygenase | -1,719592911 | 0,04127558 | 1 |
| NLA26_RS11790 |  | MerR family transcriptional regulator | 2,207672109 | 0,001235584 | 0,162433602 |
| NLA26_RS11870 |  | hypothetical protein | 1,458076504 | 0,028932455 | 1 |
| NLA26_RS11940 | cntL | D-histidine (S)-2-aminobutanoyltransferase CntL | -1,954899084 | 0,023294179 | 1 |
| NLA26_RS12135 |  | ABC transporter ATP-binding protein/permease | 1,738193519 | 0,010951952 | 0,84491089 |
| NLA26_RS12180 | hlgB | bi-component gamma-hemolysin HlgAB/HlgCB subunit B | -1,341484857 | 0,048399452 | 1 |
| NLA26_RS12190 | hlgA | bi-component gamma-hemolysin HlgAB subunit A | -1,378755834 | 0,045414938 | 1 |
| NLA26_RS12195 | sbi | immunoglobulin-binding protein Sbi | -3,226248139 | 1,06E-05 | 0,0030989 |
| NLA26_RS12210 |  | 2,3-diphosphoglycerate-dependent phosphoglycerate mutase | 2,134177701 | 0,001721042 | 0,196273659 |
| NLA26_RS12300 | nirB | nitrite reductase large subunit NirB | 1,498957138 | 0,025448806 | 1 |
| NLA26_RS12335 | nreA | nitrate respiration regulation accessory nitrate sensor NreA | 1,451777396 | 0,029655697 | 1 |
| NLA26_RS12345 | nreC | nitrate respiration regulation response regulator NreC | -1,508402286 | 0,024125403 | 1 |
| NLA26_RS12385 |  | IS1182-like element ISSau3 family transposase | -1,675320076 | 0,014161013 | 0,952418919 |
| NLA26_RS12475 |  | L-lactate permease | 1,603332658 | 0,016609813 | 1 |
| NLA26_RS12500 |  | DUF3021 domain-containing protein | 3,448503028 | 2,58E-06 | 0,000876259 |
| NLA26_RS12505 |  | LytTR family DNA-binding domain-containing protein | 4,222707423 | 2,39E-08 | 3,14E-05 |
| NLA26_RS13390 |  | lactose-specific PTS transporter subunit EIIC | -1,661919394 | 0,01675983 | 1 |
| NLA26_RS13590 | fmtB | LPXTG-anchored DUF1542 repeat protein FmtB | 1,640571816 | 0,015311673 | 1 |
| NLA26_RS13765 |  | UDP-N-acetylglucosamine 1-carboxyvinyltransferase | -2,18598313 | 0,001362386 | 0,162433602 |
| NLA26_RS14375 | tst | toxic shock syndrome toxin TSST-1 | -3,585464999 | 8,20E-07 | 0,00053791 |
| NLA26_RS14390 |  | TDT family transporter | 1,428978692 | 0,033704317 | 1 |

**Table S2.**
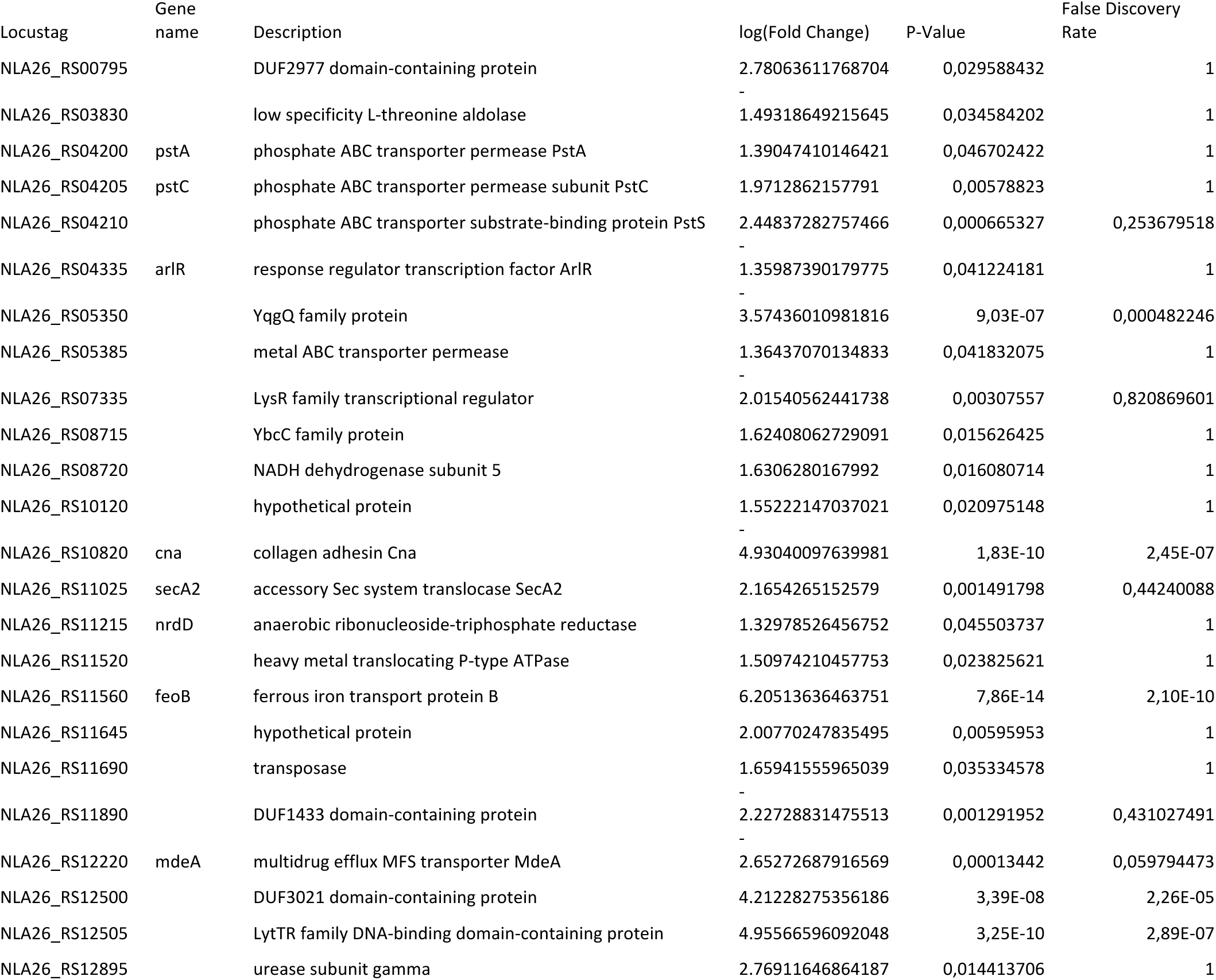
Significant transcriptional comparison analysis of all differentially expressed genes in between MN8*ΔsaeS* with or without PP-HCl.

**Table S3.**
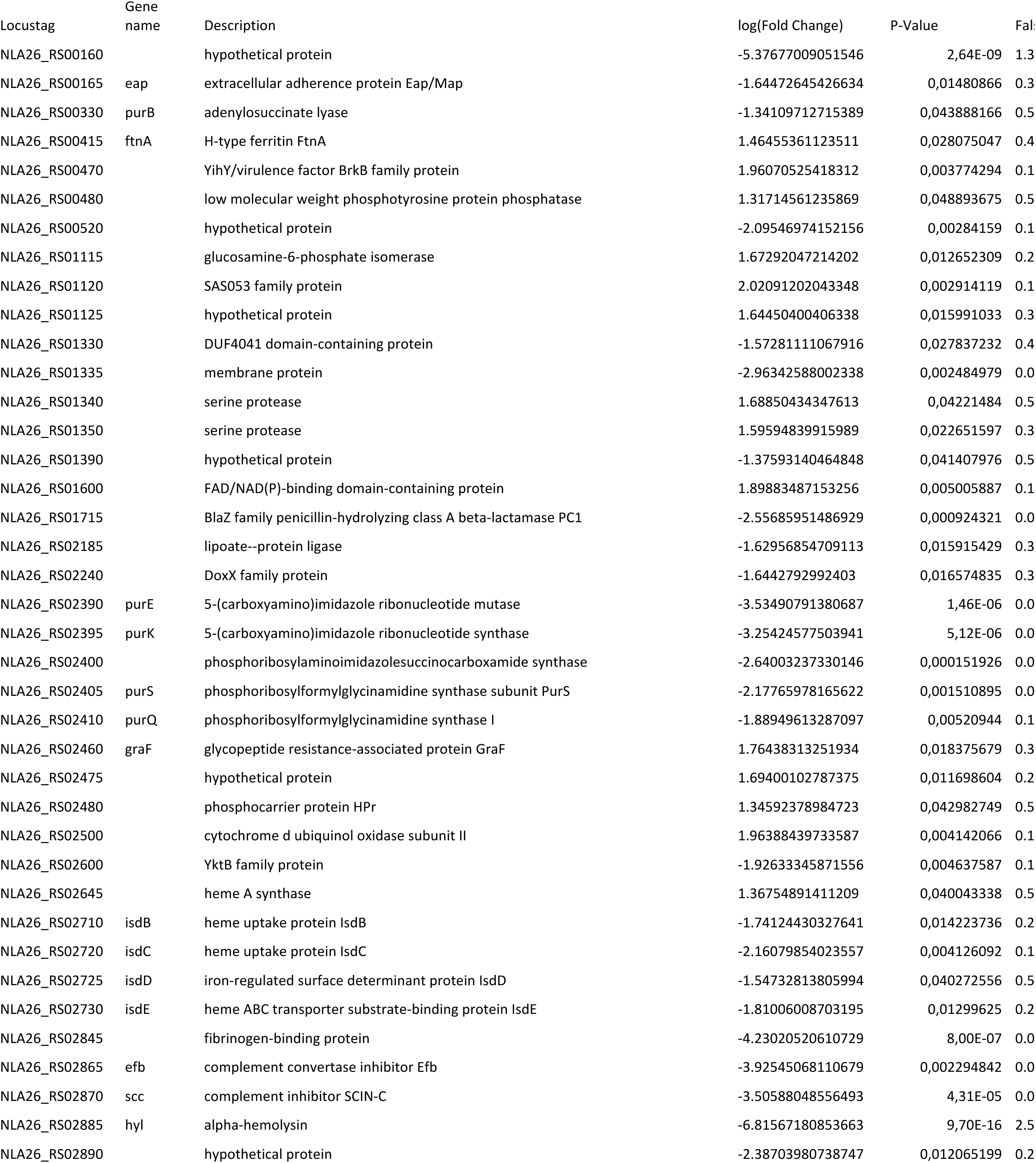
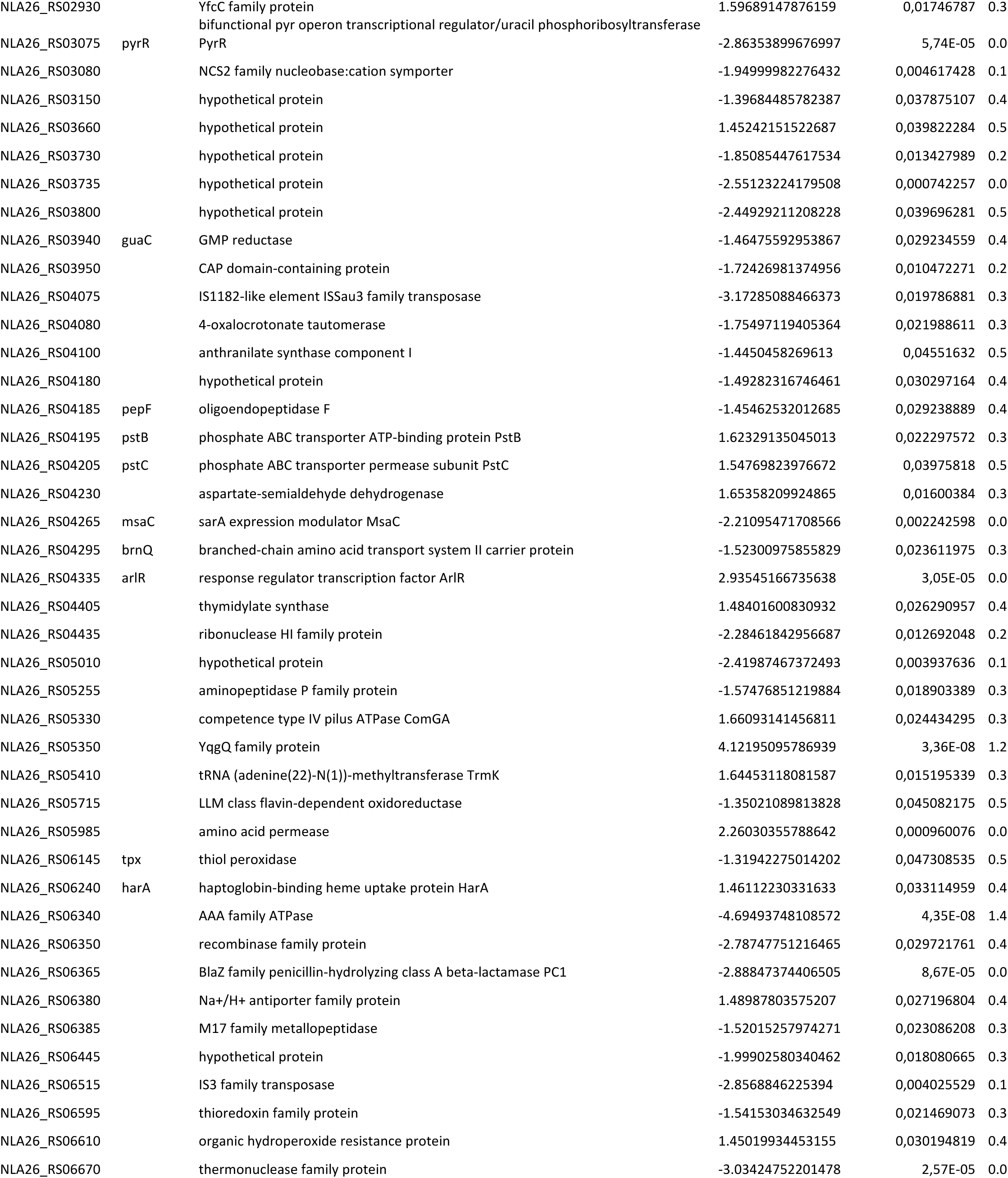
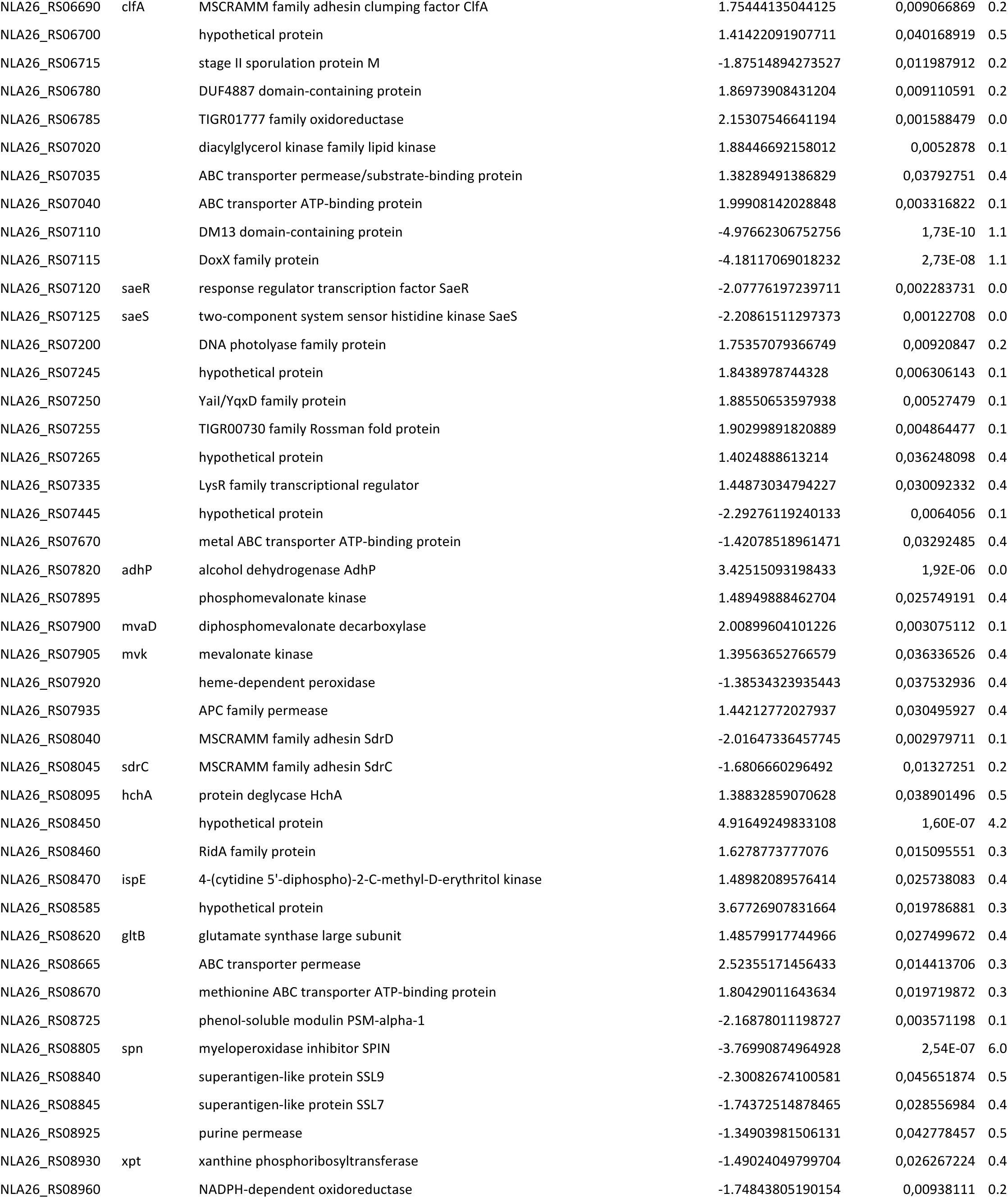
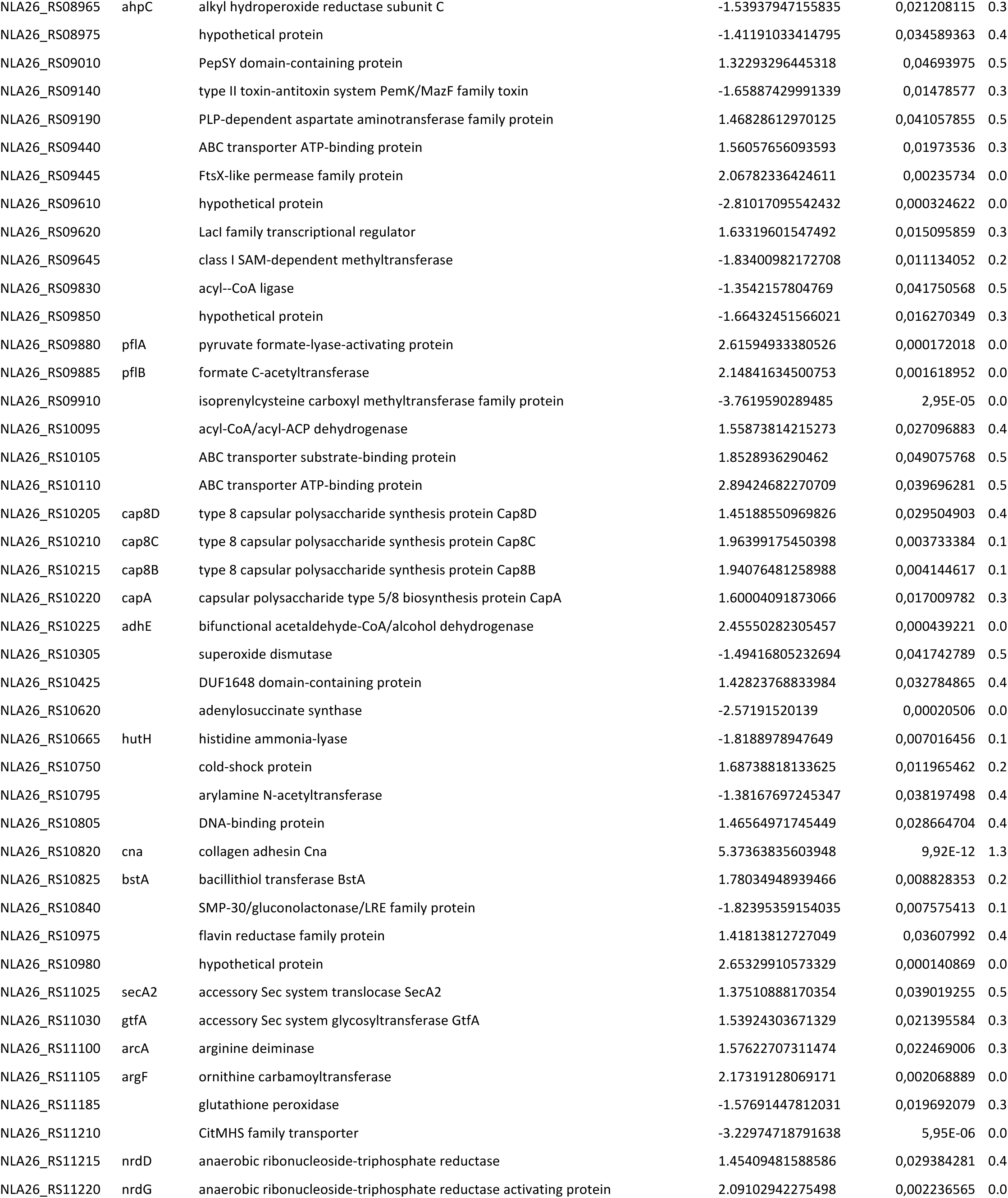
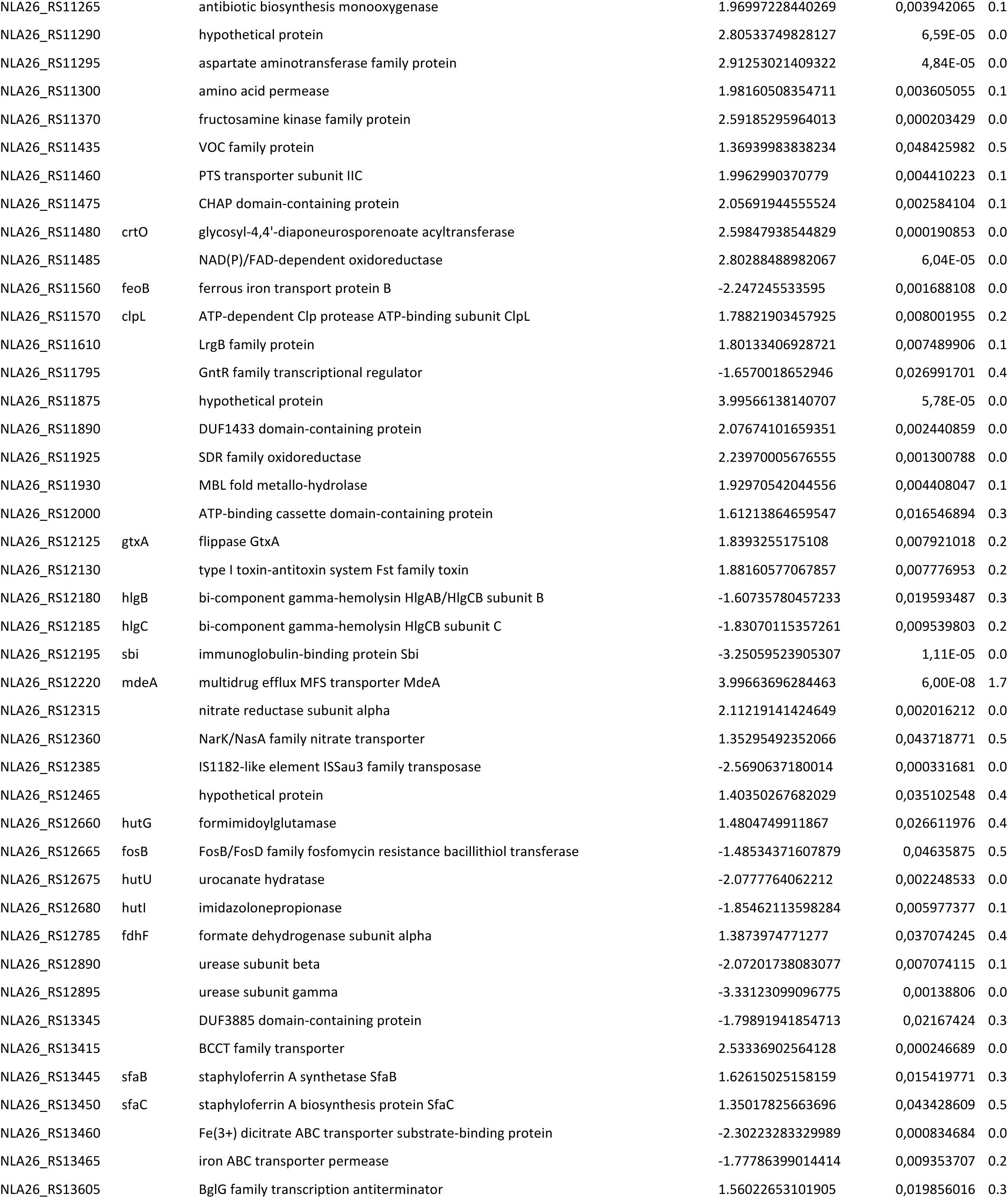
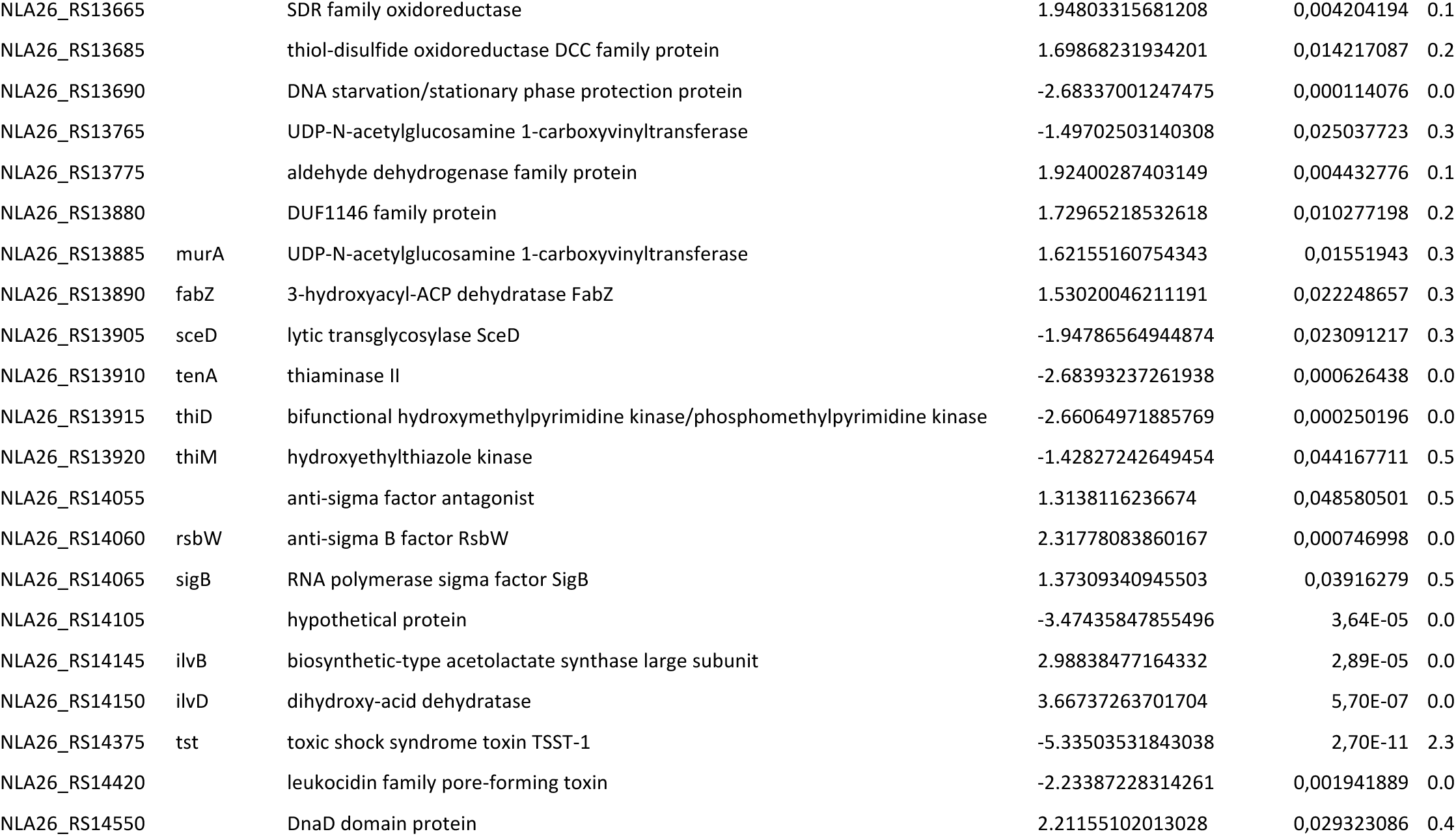
Significant transcriptional comparison analysis of all differentially expressed genes in between MN8 and MN8*ΔsaeS*.

